# Morphological Cell Profiling of SARS-CoV-2 Infection Identifies Drug Repurposing Candidates for COVID-19

**DOI:** 10.1101/2020.05.27.117184

**Authors:** Carmen Mirabelli, Jesse W. Wotring, Charles J. Zhang, Sean M. McCarty, Reid Fursmidt, Tristan Frum, Namrata S. Kadambi, Anya T. Amin, Teresa R. O’Meara, Carla D. Pretto, Jason R. Spence, Jessie Huang, Konstantinos D. Alysandratos, Darrell N. Kotton, Samuel K. Handelman, Christiane E. Wobus, Kevin J. Weatherwax, George A. Mashour, Matthew J. O’Meara, Jonathan Z. Sexton

**Author notes:** These authors contributed equally to this work (Carmen Mirabelli, Jesse W. Wotring). These authors contributed equally to this work (Charles J. Zhang, Sean M. McCarty, Reid Fursmidt). Co-Senior Authors (Matthew J. O’Meara, Jonathan Z. Sexton).

## Abstract

The global spread of the severe acute respiratory syndrome coronavirus 2 (SARS-CoV-2), and the associated disease COVID-19, requires therapeutic interventions that can be rapidly identified and translated to clinical care. Traditional drug discovery methods have a >90% failure rate and can take 10-15 years from target identification to clinical use. In contrast, drug repurposing can significantly accelerate translation. We developed a quantitative high-throughput screen to identify efficacious agents against SARS-CoV-2. From a library of 1,425 FDA-approved compounds and clinical candidates, we identified 17 dose-responsive compounds with *in vitro* antiviral efficacy in human liver Huh7 cells and confirmed antiviral efficacy in human colon carcinoma Caco-2, human prostate adenocarcinoma LNCaP, and in a physiologic relevant model of alveolar epithelial type 2 cells (iAEC2s). Additionally, we found that inhibitors of the Ras/Raf/MEK/ERK signaling pathway exacerbate SARS-CoV-2 infection *in vitro.* Notably, we discovered that lactoferrin, a glycoprotein classically found in secretory fluids, including mammalian milk, inhibits SARS-CoV-2 infection in the nanomolar range in all cell models with multiple modes of action, including blockage of virus attachment to cellular heparan sulfate and enhancement of interferon responses. Given its safety profile, lactoferrin is a readily translatable therapeutic option for the management of COVID-19.

**IMPORTANCE:** Since its emergence in China in December 2019, SARS-CoV-2 has caused a global pandemic. Repurposing of FDA-approved drugs is a promising strategy for identifying rapidly deployable treatments for COVID-19. Herein, we developed a pipeline for quantitative high-throughput image-based screening of SARS-CoV-2 infection in human cells that led to the identification of several FDA-approved drugs and clinical candidates with *in vitro* antiviral activity.

## INTRODUCTION

SARS-CoV-2 is an enveloped, positive-sense, single-stranded RNA betacoronavirus that emerged in Wuhan, China in November 2019 and rapidly developed into a global pandemic. The associated disease, COVID-19, manifests with an array of symptoms, ranging from flu-like illness and gastrointestinal distress (1, 2) to acute respiratory distress syndrome, heart arrhythmias, strokes, and death (3, 4). Recently, the FDA issued emergency approval of remdesivir, a nucleoside inhibitor prodrug developed for Ebola virus treatment (5). However, there are no established prophylactic strategies or direct antiviral treatments available to limit SARS-CoV-2 infections and to prevent the associated disease COVID-19.

Repurposing of FDA-approved drugs is a promising strategy for identifying rapidly deployable treatments for COVID-19. Benefits of repurposing include known safety profiles, robust supply chains, and a short time-frame necessary for development (6). Additionally, approved drugs can serve as chemical probes to understand the biology of viral infection and inform on the molecular targets/pathways that influence SARS-CoV-2 infection. To date, several drug repurposing screening efforts have been reported in various cell systems including non-human primate VeroE6 (7), Huh7.5 (8) and Caco-2 cells (9) with a significant overlap in reported drugs but with wide-ranging potencies. Here, we developed a pipeline for quantitative high-throughput image-based screening of SARS-CoV-2 infection that led to the identification of several FDA-approved drugs and clinical candidates with previously unreported *in vitro* antiviral activity. We also determined that inhibitors of the Ras/Raf/MEK/ERK signaling pathway exhibited proviral activity in Huh7. Mechanism of action studies of lactoferrin, the most promising hit, identified that it inhibits viral attachment, enhances antiviral host cell responses, and potentiates the effects of remdesivir.

## RESULTS

To determine the optimal cell line and assay timing for antiviral drug screening, we assessed SARS-CoV-2 infectivity in Vero E6 (African green monkey kidney cells), Caco-2 (human colon adenocarcinoma cells), Huh7 (human hepatocyte carcinoma cells) and LNCaP (human prostate adenocarcinoma). Viral growth kinetics at a multiplicity of infection (MOI) of 0.2 revealed that each cell line supported viral infection with peak viral titers at 48 hours post infection (hrs p.i.), except for Caco-2, which took 72 hrs (Fig. S1A). The Huh7 cell line was selected for drug screening because it produced the maximum percentage of infected cells (~20%) at 48 hrs p.i. at a MOI of 0.2, while Caco-2 and LNCaP required higher MOI to show the same infection rates (Fig. S1B). Huh7 also exhibited superior signal to background for N protein staining, and viral infection was detectable at an MOI of as low as 0.004 at 48 hrs p.i. (Fig. S1C).

### Cell morphological profiling of SARS-CoV-2 infected cells

To gain insight into cellular features that are being perturbed upon infection, a cell painting style morphological profiling assay was developed in 384-well plates. A multiplexed fluorescent dye set labeling the SARS-CoV-2 nucleocapsid protein (N), nuclei (Hoechst 33342), neutral lipids (HCS LipidTox Green), and cytoplasm (HCS CellMask Orange) was used to capture a wide variety of cellular features relevant to viral infectivity, including nuclear morphology, nuclear texture, and cytoplasmic and cytoskeletal features. Cell level features of infected and uninfected cells were measured using a CellProfiler (7) image analysis pipeline. We observed several prominent features associated with SARS-CoV-2 infection, including the formation of syncytia, cytoplasmic protrusions, multiple cell shapes, and positive/negative N protein staining within the nucleus. Fig. 1A shows multiplexed images of infected and uninfected wells and resulting identification/segmentation of infected cells. To systematically explore the morphologies of infected cells, features were dimensionally reduced via the non-linear uniform manifold approximation and projection (UMAP). The analysis showed five regions of interest (ROI) (Fig. 1B) with selected phenotypes. These phenotypes included rounded up cells with intense N staining overlapping with the nuclei (ROI I), diffuse N staining in the cytoplasm of cells with normal shape and size (ROI II), and cells with abnormal cytoplasmic protrusions containing punctate N staining (ROI III) or diffused N staining (ROI IV). Most infected cells, however, clustered in syncytia (ROI V), suggesting that infection in Huh7 propagates primarily through cell-to-cell fusion. Fig. 1C shows split violin plots for prominent features that are perturbed in infected vs. uninfected cells. Viral staining, cytoplasmic intensity (CellMask), and nuclear texture all increase in infected cells. In addition, the neutral lipid droplet content increases and the radial distribution of the lipid droplets shifts outwards from the nucleus towards the plasma membrane. Increased lipid accumulation has been observed previously in Hepatitis C virus-infected Huh7 cells (8). The CellMask intensity is increased in infected cells due to the prevalence of syncytia where the disappearance of cell boundaries increases staining intensity at the cell edge. Collectively, our analysis identifies specific features characteristic of SARS-CoV-2 infected cells.

**Figure 1.**
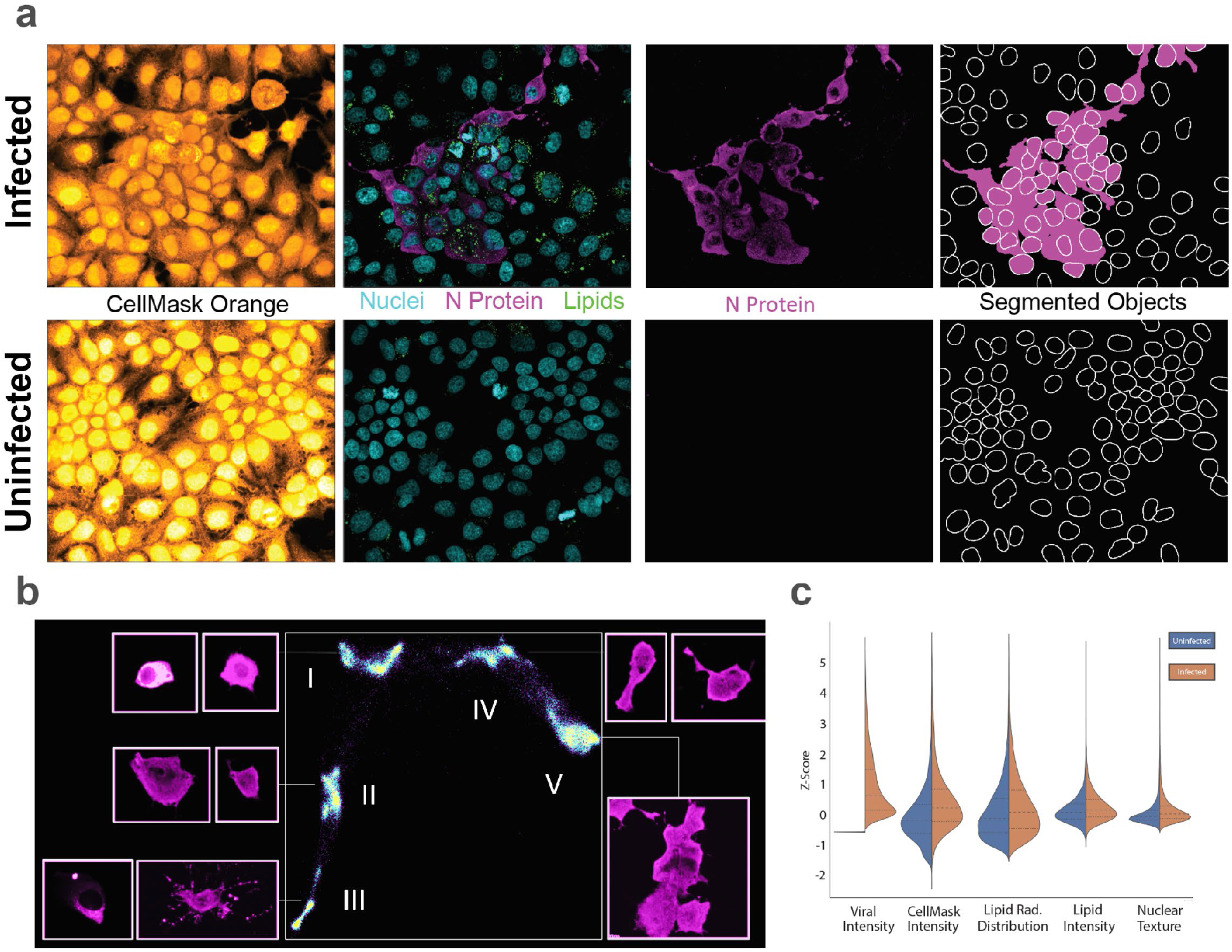
Morphological profiling of SARS-CoV-2 infected Huh7 cells (MOI of 0.2 for 48 hrs). A) Clockwise: Representative field with nuclei (cyan), neutral lipids (green), and SARS-CoV-2 N protein (magenta), N protein image in the same area with “fire” false color LUT showing distinct morphologies of infected cells showing small/round cells with a hollow center, cells with protrusions, and large syncytia, CellMask image showing cell boundaries and syncytia formation. B) UMAP embedding and phenotypic clustering of 3 million cells showing distinct morphologies present, including: small/bright cells (I), cells with protrusions (III), and syncytia (V). C) Comparison of normalized cellular features in infected (brown) and uninfected (blue) cells showing differences in cytoplasmic organization, lipid content/distribution and nuclear texture. All distributions were compared with the Mann-Whitney test and are statistically significant with P<0.0001.

### Identification of FDA-approved drugs with antiviral activity against SARS-CoV-2

To identify compounds with antiviral activity against SARS-CoV-2, we tested a library of 1,425 FDA-approved compounds and rationally included clinical candidates (Supplementary File 1) in Huh7 cells in quantitative high-throughput screening (qHTS) at five concentrations (50 nM, 250 nM, 500 nM, 1000 nM and 2000 nM). Compounds were assessed for their antiviral activity (shown schematically in Fig. 2A) using a CellProfiler (7) image analysis pipeline to: 1) identify infected objects in the N protein image (from a single cell to large syncytia), 2) measure their morphologic features, and then 3) tabulate how many nuclei reside within the infected objects to calculate the total percentage of infected cells per well. To increase the accuracy of identifying true actives and decrease the false negative rate of the assay, a liberal selection criteria was employed to choose drugs for follow-up studies (see methods and Fig. 2A). 132 drugs were selected from qHTS screening or by known activity against SARS, MERS or SARS-CoV-2 and carried forward for triplicate dose-response confirmation. Ultimately, 17 dose-responsive compounds were confirmed with IC_50_ values of less than 1 μM (Fig. 2B and Supplementary Table 1). These compounds include ten that are novel *in vitro* observations (domperidone, entecavir, fedratinib, ipratropium bromide, lactoferrin, lomitapide, metoclopramide, S1RA, thioguanine, and Z-FA-FMK), and seven previously reported (amiodarone, bosutinib, verapamil, gilteritinib, clofazimine (9, 10), niclosamide (11), and Remdesivir(12)). The remaining compounds either lacked efficacy, exhibited cytotoxicity (e.g. Digoxin), or were efficacious only at concentrations above 1 μM (e.g. hydroxychloroquine, chloroquine) and were thus not prioritized for follow-up. Collectively, the 17 identified hits could be stratified by compound class as ion channel modulators (amiodarone, verapamil, clofazimine, and S1RA), nucleosides/DNA binders (remdesivir, entecavir, niclosamide, and thioguanine), kinase inhibitors (bosutinib, fedratinib, and gilterinib) and others (Table 1).

**Figure 2.**
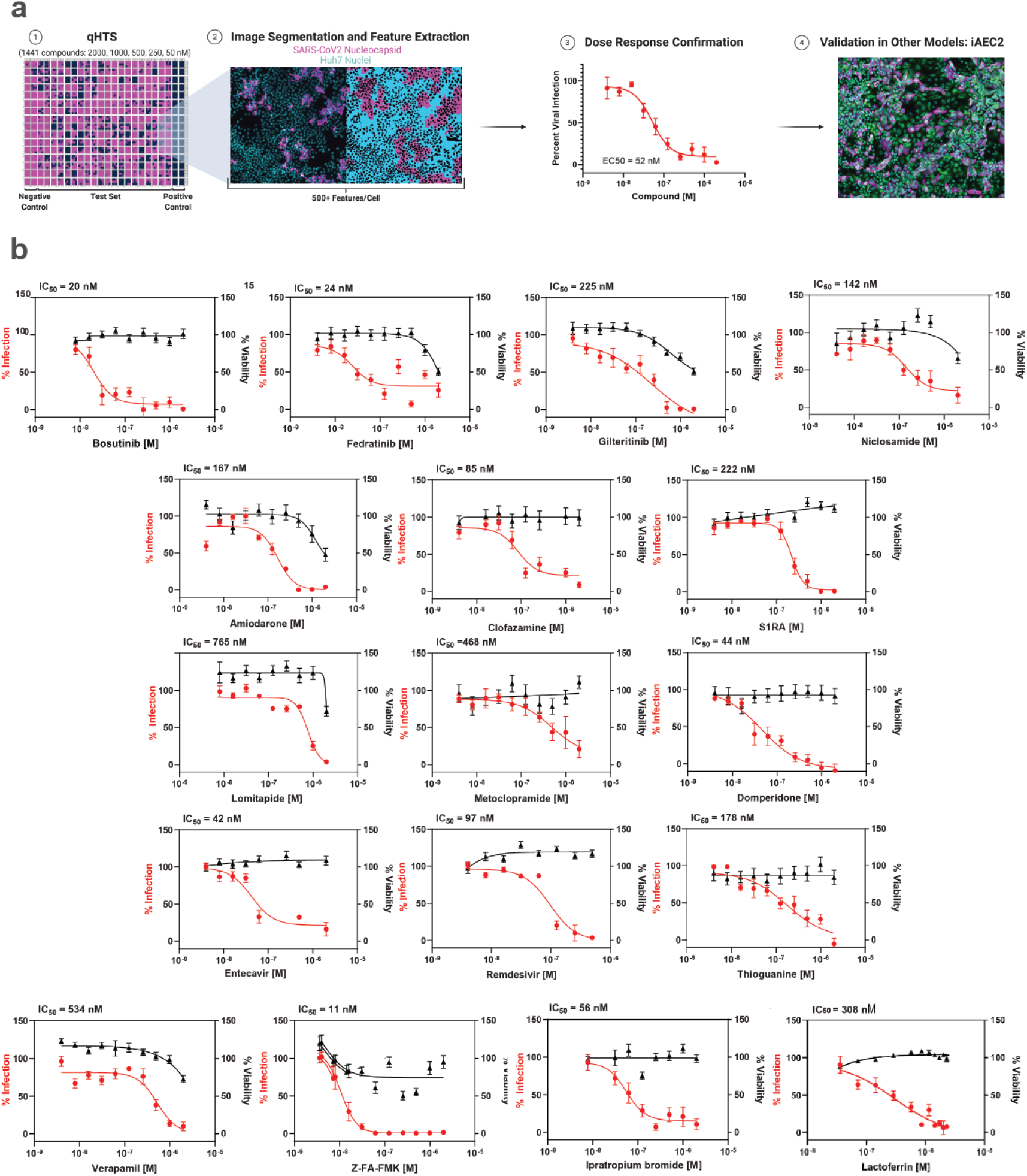
A) Schematic of the anti-SARS-CoV-2 drug repurposing screening. 1) Compounds are administered in qHTS to cells cultured on 384-well plates infected with SARS-CoV-2 and incubated for 48-hours. Each plate contains 32 negative (infected) and 32 positive (non-infected) control wells. 2) Cells are fixed, stained, and imaged. Images are analyzed using Cell Profiler to identify nuclei, cell boundaries, neutral lipid content and viral staining intensity. 3) Dose-response curves are fit to normalized percent infected cells per well. 4) Confirmation of antiviral activity in other cell lines, including a physiological relevant iPSC-derived human alveolar epithelial cell (iAECs); B) Dose-response curves of 17 compounds. Graphs represent median SEM of 10-point 1:2 dilution series of selected compounds for N=3 biological replicates. IC50 values were calculated based on normalization to the control and after fitting in GraphPad Prism.

**Table 1:** Compound Summary

| ID Drug Bank ID | Compound name | Indication | General mechanism of action | Potential mode of action against SARS-CoV2 | IC50 (nM) Huh-7 | IC50 (nM) Vero E6 | IC50 (nM) Caco-2 E6 | IC50 (nM) LNcaP E6 | IC50 (nM) iAEC2 |
| --- | --- | --- | --- | --- | --- | --- | --- | --- | --- |
| DB01118 | Amiodarone (hydrochloride) | Treatment of ventricular tachycardia | Inhibitor of K and Ca channels | Entry inhibitor<br>Evaluated in clinical trial for COVID19 (NCT04351763) | 167 | 1406 | 5500 | >5000 | 118 |
| DB06616 | Bosutinib | Treatment of chronic myeloid leukemia | Bcr-Abl kinase inhibitor | Inhibitor of S protein fusion similar to the related Imatinib | 20 | >5000 | IA | Inverse | IA |
| DB00845 | Clofazimine | Treatment of leprosy | Binds to mycobacterial DNA and K transporters inhibitor | Entry inhibitor<br>Evaluated in clinical trial for COVID19 (NCT04465695) | 85 | >5000 | >5000 | 29 | >5000 |
| DB01184 | Domperidone | Antiemetic | Dopamine D2 receptor antagonist | Host modulation | 44 | IA | IA | ND | IA |
| DB00442 | Entecavir (hydrate) | Treatment of hepatitis B virus | Transcription inhibitor, nucleoside analog | Replication inhibitor | 42 | >5000 | IA | IA | IA |
| DB12500 | Fedratinib | Treatment of intermediate-2 and high risk primary and secondary myelofibrosis | Tyrosine kinase inhibitor (Jak1) | Predicted inhibitor of kinase (NAK) family reported to reduce viral infection in vitro. Reduction of TH17 responses responsible of SARS-CoV-2 associated cytokine storm | 24 | >5000 | >5000 | >5000 | 1810 |
| DB12141 | Gilteritinib | Treatment of FLT3-mutated acute myeloid leukemia (AML) | FMS-like tyrosine kinase 3 (FLT3) inhibitor | Host modulation | 225 | 2344 | IA | IA | >5000 |
| DB00332 | Ipratropium Bromide | Treatment of COPD and asthma | Muscarinic receptor antagonist | Binding inhibitor | 56 | NA | 85 | >5000 | 4 |
| NA | Lactoferrin | Diet supplement | Iron chelator, immune-modulator<br>Antimicrobial activity | Entry and post-entry inhibitor | 308 | NA | 1170 | 157 | 45 |
| DB08827 | Lomitapide | Treatment of homozygous familial hypercholesterolemia | Microsomal triglyceride transfer protein (MTP) inhibitor | Host lipid metabolism modulation | 765 | 1875 | >5000 | IA | 733 |
| DB01233 | Metoclopramide | Treatment of diabetic gastroparesis | Dopamine D2 and serotonin 5-HT3 receptor inhibitor | Binding inhibitor | 468 | >5000 | IA | IA | IA |
| DB06803 | Nicosamide | Treatment of tapeworm infections | Mitochondrial uncoupler, mTORC1 inhibitor | Autophagy and endocytic pathway inhibitor | 142 | >5000 | IA | >5000 | NA |
| DB14761 | Remdesivir | Investigational for Ebola virus treatment | Transcription inhibitor, nucleoside analog | Replication inhibitor<br>Emergency FDA approval for COVID19 | 97 | NA | 19 | 106 | 18 |
| ZINC95000617 | S1RA | Treatment of neuropathic pain (phase II) | Sigma R1/R2 modulator | Entry inhibitor | 222 | >5000 | >5000 | IA | 1 |
| DB00352 | Thioguanine | Therapy of acute leukemia | Guanine analogue | Replication inhibitor | 178 | 820 | 674 | 29 | IA |
| DB00661 | Verapamil (hydrochloride) | Treatment of high blood pressure, heart arrhythmias, and angina | Ca channel inhibitor | Entry inhibitor<br>Evaluated in clinical trial for COVID19 (NCT04351763) | 534 | >5000 | Low | 18 | NA |
| CAS 197855-65-5 | Z-FA-FMK | Preclinical | Irreversible inhibitor of cysteine proteases (cathepsin L) | Entry inhibitor | 11 | 51 | Low | 514 | 4 |

NA=not tested, IA-inactive, >5000 indicates activity but loss of potency, Low=IC50 below 1nM. Grey rows indicate compounds with efficacy across multiple cell systems.

### Hit validation in Caco-2, LNCaP, Vero E6 and an iPSC-derived model of alveolar epithelial cells, the iAEC2

To evaluate the translatability of the 17 hits from Huh7 cells in other cell systems, we confirmed activity in LNCaP, Caco-2, and Vero E6 cell lines and in physiologically-relevant iPSC-derived alveolar epithelial type 2 cells (iAEC2s) (13). Antiviral activities across the cell systems are shown in Table 1. iAEC2s were used as a biomimetic model of the human bronchial epithelium that is involved in COVID-19 pathogenesis (14). iAEC2s are permissive to SARS-CoV-2 infection, exhibiting 10-20% N protein-positive cells at MOI of 0.2 and 50-60% positivity at MOI of 10. Upon infection, we observed long tubular protrusions that co-stained with viral N protein (Fig. 3A). Additionally, unlike the Huh7 model, the vast majority of infected iAEC2 cells were not present in viral syncytia, suggesting that cell-to-cell spread by cell fusion is limited in this model. Nine out of the 17 hits - amiodarone, lomitapide, ipratropium bromide, gilteritinib, fedratinib, and clofazimine, remdesivir, S1RA and bovine lactoferrin-showed dose-responsive antiviral activity against SARS-CoV-2 in iAECs (Table 1). Remarkably, even at a high MOI of 10, bovine lactoferrin, human lactoferrin, S1RA, and remdesivir (IC_50_ = 18 nM) retained antiviral activity, reflecting the strong efficacy of these compounds in virus-saturated infection conditions (Fig. 3B). Six compounds (Amiodarone, Ipatropium Bromide, Lactoferrin, Lomitapide, Remdesivir, Z-FA-FMK) maintained efficacy across all tested cell systems (Table 1), suggesting the targets are conserved across multiple cell types.

**Figure 3.**
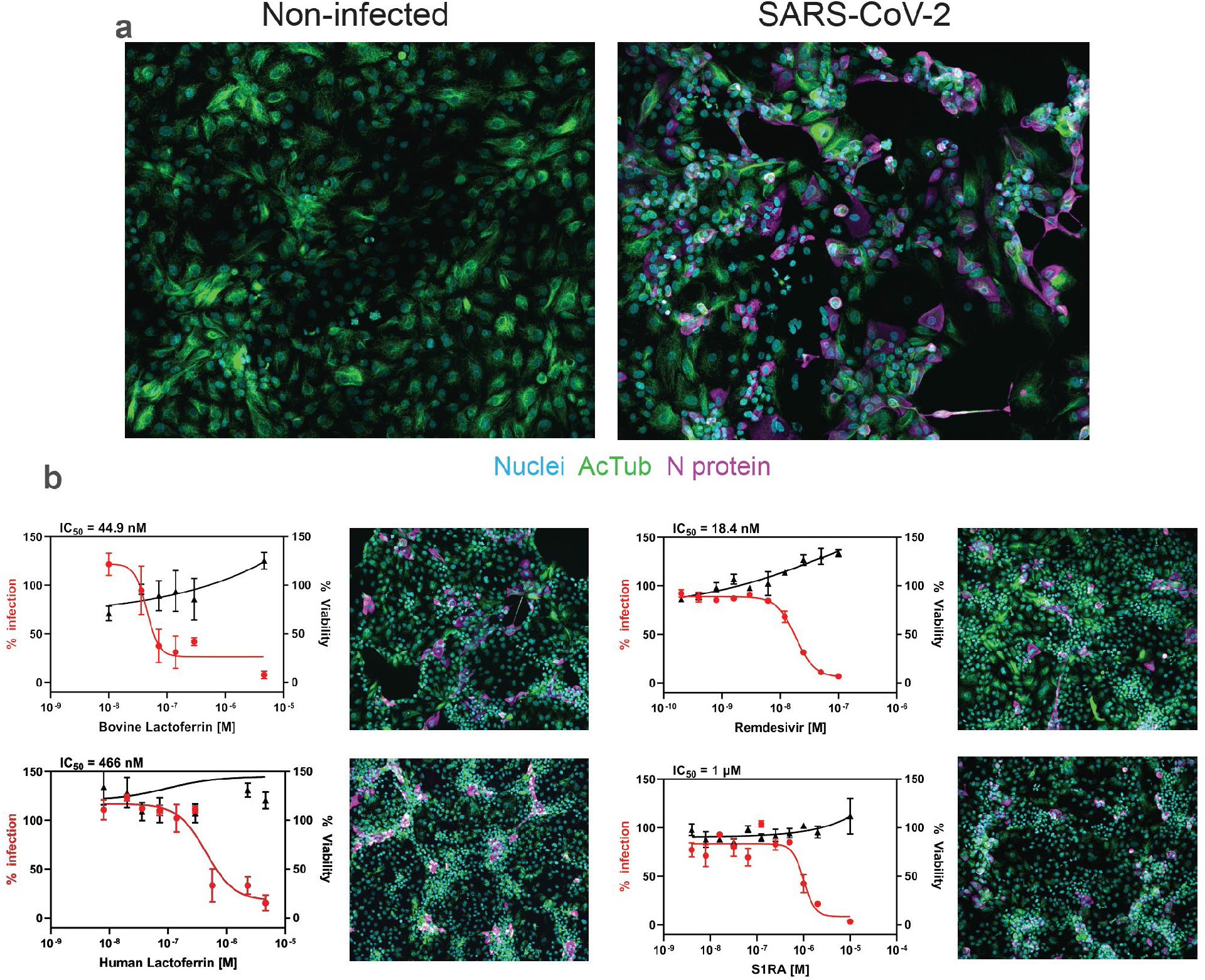
A) SARS-CoV-2 infected iAEC2 cells at MOI of 10, 48 hrs p.i. Nuclei are in cyan, N protein in magenta, and acetylated tubulin in green. Representative image was acquired on a Yokogawa CQ1 high-content imager with a 60X lens and visualized with Fiji ImageJ. B) Antiviral activity of bovine and human lactoferrin, remdesivir, and S1RA was assessed in iAEC2 cells infected with SARS-CoV-2 at MOI 10. Graphs represent median SEM of 10-point 1:2 dilution series of selected compounds for N=3 biological replicates. Representative Images of nuclei (cyan), acetylated tubulin (green), and N protein (magenta) at compound IC_50_ dose are also represented.

### Characterization of antiviral hits and identification of compounds that exacerbate viral infection

To stratify compounds, we performed a time-of-addition study with compound added either 4 hrs prior to infection (as done previously in the screen) or 1 hr p.i. (Fig. 4A). We infected Huh7 with SARS-CoV-2 at MOI of 1 and then quantified infection by detecting the positive-strand viral RNA genome by RNAscope (Fig 4B). We found that verapamil, entecavir, and niclosamide lost activity under these experimental conditions (Fig. 4C). Amiodarone, clofazimine, S1RA, lomitapide, Z-FA-FMK, the other nucleoside analogues remdesivir and thioguanine, and the kinase inhibitors bosutinib, fedratinib, and gilterinib retained activity regardless of compound addition pre- or post-infection, suggesting that they inhibit post-binding entry events. Two compounds, ipratropium bromide and metoclopramide, lost activity when added 1 hr p.i., suggesting a role in viral binding inhibition. Although they share the same molecular target (dopamine D2 receptor), metoclopramide and domperidone seem to exert their antiviral activity with different modes of action, either by directly inhibiting binding or indirect effects on the host.

**Figure 4.**
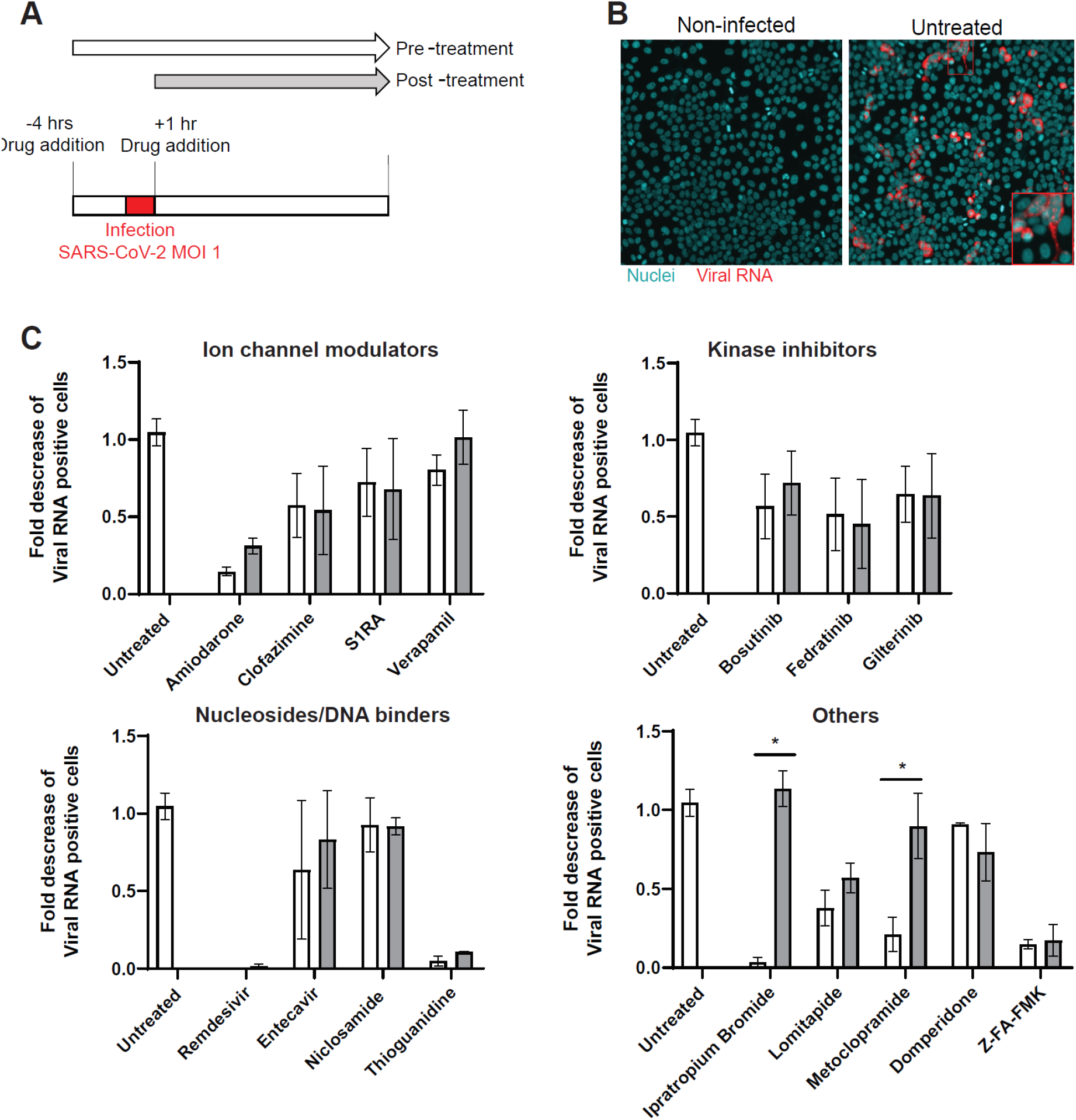
Time of drug-addition of the identified antiviral hits. A) Experimental scheme where compounds are added 4 hrs prior (same treatment window as drug screening) or 1 hr postinfection (p.i.) with SARS-CoV-2 (MOI of 1). Huh7 cells are fixed, permeabilized and subjected to RNAscope analysis 48 hrs p.i. B) Representative image of SARS-CoV-2 infected and non-infected Huh7 cells acquired on the CX5 high-content platform at 10X and analyzed with Fiji ImageJ. Viral RNA is represented in red, nuclei in cyan. C) Time of drug-addition for selected antiviral hits (at 10x IC_50_ dose) organized according to the compound class. Graphs represent the fold decrease of infection over the untreated condition. Infection was calculated on the viral RNA image after image segmentation with Cell Profiler. Graphs represent an average SEM of N=3 biological replicates. Statistical significance determined using multiple student’s t-test with the Bonferroni-Dunn correction method, with alpha = 0.05. *p<0.01.

Our screening also identified compounds that exacerbated infection. All Mitogen/Extracellular signal-regulated Kinase (MEK) inhibitors tested (cobimetinib, trametinib, and binimetinib) resulted in a >2-fold increase of viral infection in Huh7 (Fig. 5A, B). To confirm this finding, we performed RNAscope on virus-infected, cobimetinib-treated versus untreated cells 24 and 48 hrs p.i. (Fig. 5C). The percentage of viral RNA-positive cells was increased at 48 hrs p.i., but not at 24 hrs p.i., following treatment, suggesting that these compounds could enhance virus spread. In addition, upon treatment with the three MEK inhibitors, and cobimetinib in particular, we observed an increased syncytia size (Fig 5A) and different patterns of localization of viral RNA and S protein within the infected cells (Fig 5C). These immunofluorescence staining patterns suggest a difference in viral compartmentalization and spread in MEK inhibitor-treated cells. The increased infection and the diffuse localization of viral RNA was recapitulated when treating the cells with a molecular probe, U0126 (10 μM), that is commonly used as an inhibitor of the Ras-Raf-MEK-ERK pathway. Taken together, these data highlight the utility of screening FDA-approved compounds as a way of identifying cellular pathways involved in viral infection.

**Figure 5.**
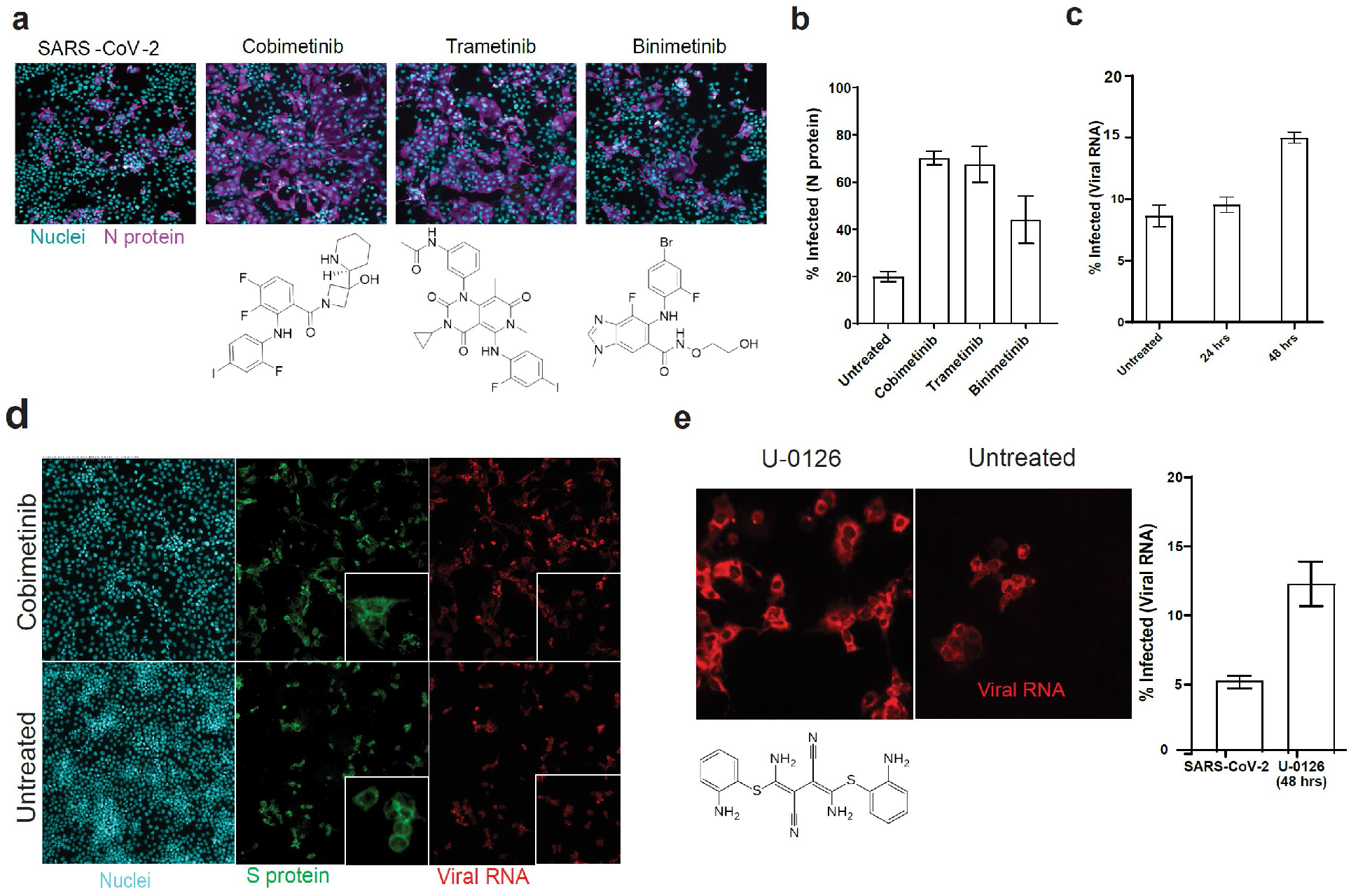
Selective MEK inhibitors exacerbate SARS-CoV-2 infection. A) Representative images of Huh7 cells infected with SARS-CoV-2 (MOI of 0.2) and treated with cobimetinib (250 nM), trametinib (250 nM), and binimetinib (250 nM) with nuclei in cyan and N protein in magenta. Viral infection was calculated on N protein images after image segmentation with Cell Profiler. Bars represent N=3 technical replicates and unpaired t-tests with Welch’s correction were performed in Graphpad Prism. * p<0.001. B) RNAscope of Huh7 infected with SARS-CoV-2 (MOI of 1) treated with cobimetinib (1000 nM) and harvested at 24 hrs and 48 hrs p.i. Graph represents average, SEM of N=3 biological replicates. D) Representative images of SARS-CoV-2-infected (MOI of 1) and cobimetinib (1000 nM)-treated Huh7. Cells were harvested 48 hrs p.i., subjected to RNAscope to detect viral RNA (positive strand, in red) and counterstained with anti-S protein antibody (green) and Hoechst 33342 (nuclei in cyan). E) SARS-CoV-2-infected (MOI of 1) Huh7 were treated with U-0126 (10 μM) and subjected to RNAscope 48 hrs p.i. Graph represents average SEM of N=2 biological replicates, each with three technical replicates.

### Lactoferrin blocks SARS-CoV-2 replication at different stages of the viral cycle

The most broadly efficacious hit identified was lactoferrin, a protein found in colostrum and airway epithelium (15). To confirm our previous finding of inhibition of N protein expression by lactoferrin, we infected Huh7 cells with SARS-CoV-2 (MOI of 0.2) under increasing doses of lactoferrin and measured viral RNA using RT-qPCR at 48 hrs p.i. (Fig. 6A). Lactoferrin exhibited a dose-dependent inhibition of viral replication (Fig. 6A) and retained antiviral activity through a range of MOIs (Fig. 6B). It maintained antiviral activity even when added 1 or 24 hrs after infection, suggesting multiple modes of antiviral action (Fig. 6B). Previous work on lactoferrin in the context of SARS-CoV-1 suggested that lactoferrin blocks viral entry by binding heparan sulfate proteoglycans, which are viral attachment factors (16). Therefore, we performed a viral binding assay by incubating SARS-CoV-2 (MOI of 10) in the presence of lactoferrin (1250 nM and 6250 nM) for one hour on ice followed by quantification of viral RNA by RT-qPCR (Fig. 6C). Remdesivir was included as a negative control as it blocks viral infection at a post-binding step. Both concentrations of lactoferrin, but not remdesivir, blocked SARS-CoV-2 attachment to Huh7 cells (Fig. 6C). As a positive control, Huh7 cells were treated with NaClO_3_, a protein sulfation inhibitor that depletes cells of heparan sulfate (17). SARS-CoV-2 binding to cells was reduced in NaClO_3_-treated cells, and additional lactoferrin treatment did not further reduce binding (Fig. 6C). These data suggest that, similar to SARS-CoV-1, lactoferrin blocks viral attachment via neutralizing heparan sulfate proteoglycans. Another potential mechanism of action of lactoferrin is through enhancement of interferon responses, which can then limit viral replication within host cells (18). We therefore evaluated mRNA levels of IFNβ and the interferon-stimulated genes ISG15, MX1, Viperin and IFITM3 in lactoferrin-treated Huh7 cells (Fig. 6D). SARS-CoV-2 infection did not result in a robust interferon response consistent with previous studies (19). However, we did detect an upregulation of IFNβ and ISG transcripts in virus-infected and lactoferrin-treated cells, suggesting that the post-entry antiviral activity of lactoferrin is interferon-mediated. To rule out iron chelation as a potential mode of action, iron-saturated hololactoferrin and transferrin were tested in Huh7 cells; the former retained activity and the latter was inactive (Fig. 6E). Given the pronounced single-agent efficacy of lactoferrin, we further tested whether combinations with the FDA-approved agent remdesivir could improve the overall antiviral activity. We found that lactoferrin potentiated the efficacy of remdesivir 8-fold (Fig. 6F), suggesting that lactoferrin could be beneficial in the management of COVID-19 both as a single agent or in combination.

**Figure 6.**
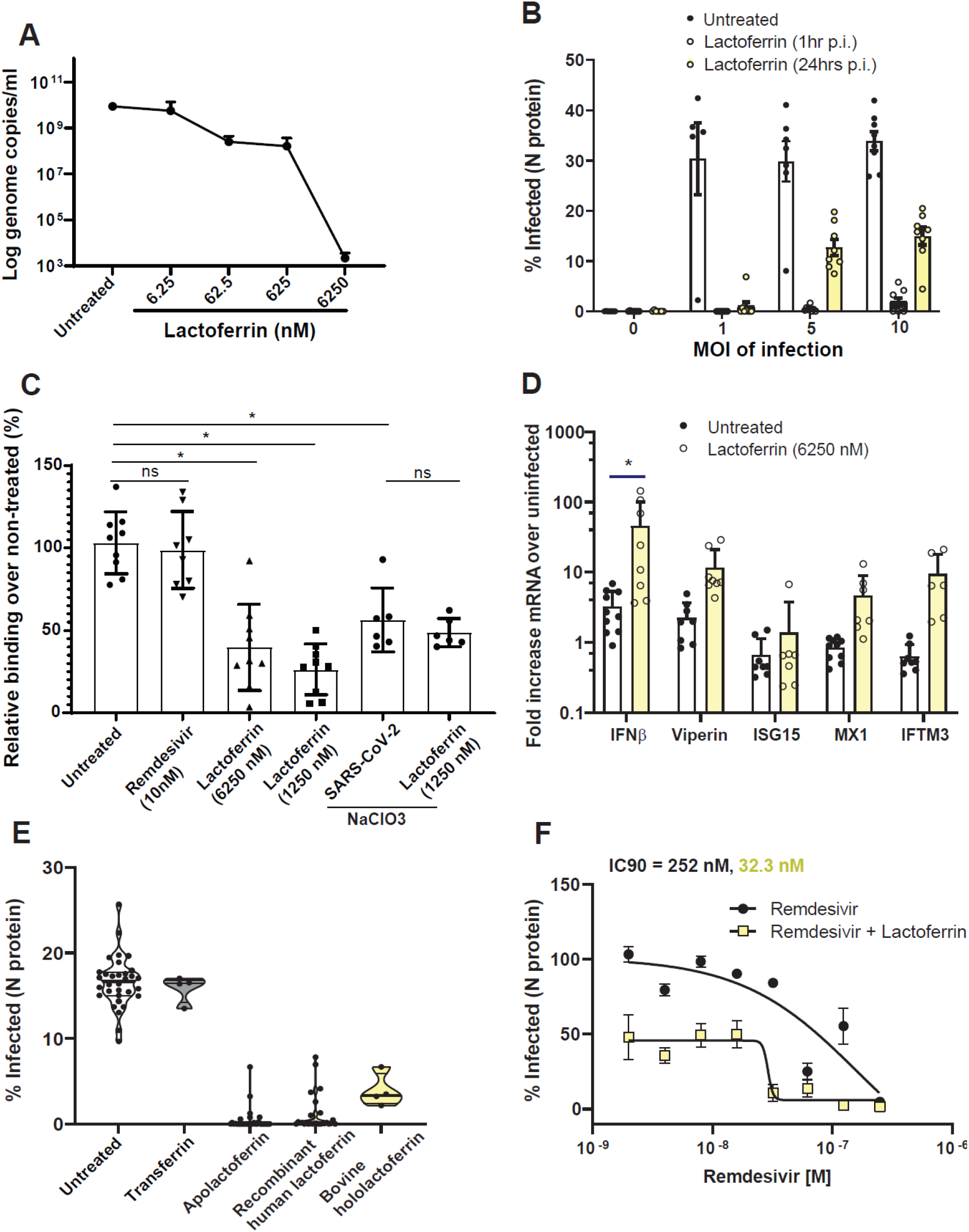
Lactoferrin blocks SARS-CoV-2 replication at different stages of the viral cycle. A) Huh7 cells were infected with SARS-CoV-2 at MOI of 0.2 for 48 hrs and treated with increasing concentration of lactoferrin (5.75 nM – 5750 nM). Cells were harvested and RNA was extracted. Viral genome copies were calculated by RT-qPCR with an absolute quantification method. B) Huh7 were infected with SARS-CoV-2 (MOI of 1, 5 and 10; MOI of 0 indicates non-infected cells) and treated with 2.3 μM of lactoferrin at 1 and 24 hrs p.i. Bars indicate the percentage of infected cells in different conditions. Data is an average of eight replicates. Statistical significance determined using multiple student’s t-test with the Bonferroni-Dunn method, with alpha = 0.05. Except for MOI of 0, all conditions (Untreated vs Lactoferrin, 1 hr or Untreated vs Lactoferrin, 24 hr) differ by p<0.0001. C) Binding assay. Huh7 cells were pre-incubated on ice with compounds: lactoferrin (100 μg/mL and 500 μg/mL) and remdesivir (10 nM), as a negative control, for one hour and then infected with SARS-CoV-2 (MOI of 10) for 1h on ice. Cells were then washed thoroughly with PBS to remove unbound virus and viral RNA was quantified by RT-qPCR. Huh7 cultured in NaClO_3_ for 7 days, which strips heparan sulfate proteoglycans from the cell surface, were used as a control for lactoferrin mode of action. Data is shown after normalization to viral control (100%) and represent an average of N=3 biological replicates with N=2-3 technical replicates each. Unpaired t-tests with Welch’s correction were performed in Graphpad Prism to determine significance. *p<0.0001. D) mRNA levels of cellular IFNβ, MX1, ISG15 and IFITM3 were calculated with ΔΔCt in SARS-CoV-2 infected and lactoferrin (5750 nM)-treated cells over uninfected Huh7. Data are average SD of N=2 biological replicates with n=3 technical replicates each. Statistical significance was determined using multiple student’s t-test with the Bonferroni-Dunn method with alpha = 0.05. *p<0.001. E) Percentage of SARS-CoV-2 infected Huh7 cells upon treatment with bovine apolactoferrin and hololactoferrin, native human lactoferrin, and transferrin at a concentration of 2.3 μM. F) Dose response of lactoferrin (0 to 2.3 μM) in combination with remdesivir (0 to 30 nM). Cells were pre-treated with combination or single therapy and infected with SARS-CoV-2 (MOI of 0.2) for 48 hrs.

## DISCUSSION

In this study, we developed an experimental workflow based on high-content imaging and morphological profiling that allows for rapid screening of FDA-approved compounds, and identified 17 compounds that inhibit SARS-CoV-2 infection *in vitro*. Of these, seven were previously reported and serve as a benchmark validation of our endpoints and experimental approach, and ten were hitherto unknown. We evaluated the antiviral activity of the 17 hits identified in Huh7 in three transformed cell lines (VeroE6, Caco-2, and LNcaP) and one non-transformed cell line (iAECs) and observed six compounds (amiodarone, ipratropium bromide, lactoferrin, lomitapide, remdesivir, Z-FA-FMK) exhibiting activity across multiple cell lines.

Since the completion of this study (June 2020), over 30 studies reporting SARS-CoV-2 antiviral activity of FDA-approved drugs have been published. A meta-analysis of these *in vitro* screens (including this effort) show consensus around 11 compounds, with small total overlap between studies (20). This observation suggests that drug screening of FDA-approved compounds is highly dependent on the chosen cell line and infection conditions. It is expected that compounds exerting an antiviral effect through direct binding to viral proteins would be more independent of the chosen cell system rather than drugs modulating host cell factors that can vary widely. For example, we observed conservation of activity across cell systems for remdesivir, which directly inhibits the viral polymerase (21), and also for lomitapide, which is proposed to directly inhibit recombinant SARS-CoV-2 main protease (Mpro)(22).

As most FDA-approved drugs are optimized against human molecular targets, active compounds can lead to target identification of host factors involved in SARS-CoV-2 infection. Z-FA-FMK is an irreversible cathepsin L inhibitor that exhibits potent antiviral activity in all of the five cell systems tested herein, presumably because cathepsin L has been shown to be an entry factor of SARS-CoV-2 (23, 24). Another hit in our Huh7 screen, fedratinib, was approved by the FDA in 2019 for myeloproliferative neoplasms (25) and is an orally bioavailable semi-selective JAK1/JAK2 inhibitor. JAK inhibitors have been proposed for COVID-19 treatment to specifically inhibit Th17-mediated inflammatory response (26, 27) and to block numb-associated kinase responsible for clathrin-mediated viral endocytosis (28). The JAK inhibitor baricitinib (29) in combination with remdesivir was recently granted emergency use authorization by the FDA, while jakotinib (<u>ChiCTR2000030170</u>), and ruxolitinib (<u>ChiCTR2000029580</u>) are currently being evaluated in clinical trials for COVID-19 as potential dual action therapeutics (antiviral and innate immune response).

In contrast to the antiviral drug hits, all FDA-approved MEK inhibitors tested exacerbated viral infection, likely by increasing cell-to-cell spread as suggested by the formation of larger syncytia and more diffuse localization of viral RNA and S protein within infected cells (Fig. 5). Intriguingly, in the context of other virus infections, including SARS-CoV-1, pharmacological inhibition of the Ras-Raf-MEK-ERK pathway results in restriction of viral infection (30). This underscores the importance of this pathway during viral infections and warrants further examination into the mechanism of action of this signaling cascade during SARS-CoV-2 infection.

This study has generated several clinically testable and readily translatable hypotheses. As an example, we observed potent antiviral activity of ipratropium bromide (Atrovent), a quaternary ammonium salt and muscarinic receptor antagonist that is commonly prescribed for asthma. It is administered via inhalation into the lungs with little systemic absorption. Given its potential mode of action as inhibitor of SARS-CoV-2 attachment, prophylaxis or post-exposure treatment with ipratropium bromide may curb infection of the upper respiratory tract and drastically reduce systemic viral spread and development of severe symptoms while achieving beneficial bronchodilation. Similarly, we identified metoclopramide and domperidone, both dopamine D2 receptor antagonists used to treat gastrointestinal symptoms, as SARS-CoV-2 inhibitors. Gastrointestinal symptoms have been increasingly reported in more than half of the patients infected by SARS-CoV-2 (2). Hence, these compounds may ameliorate GI symptoms during COVID-19 infection, and in addition the reduced viral load in the GI tract could also reduce fecal-oral transmission of SARS-CoV-2 (31).

Most noteworthy, our screen identified bovine lactoferrin, a safe and widely available dietary supplement, with multi-modal efficacy in multiple cell systems, including non-transformed and physiologically relevant iAEC2s. Lactoferrin gene expression was shown to be highly upregulated in response to SARS-CoV-1 infection (32) and, in addition to enhancing natural killer cell and neutrophil activity, lactoferrin blocks SARS-CoV-1 attachment through binding to heparan sulfate proteoglycans (15). Here, we showed that lactoferrin has a multi-modal mechanism of action against SARS-CoV-2 infection (Fig. 6). First, it strongly inhibits cellular binding of SARS-CoV-2 to cells via competition with heparan sulfate. Second, it retains antiviral efficacy at 24 hrs p.i. and it modulates host cell immunity through increased expression of interferon and several interferon-stimulated genes. Through heightening the innate immune response of host cells, orally administered lactoferrin could be especially effective in resolving the gastrointestinal symptoms that are present in COVID-19 patients (33) with a mechanism similar to norovirus infection (34). In addition, lactoferrin was previously shown to decrease the production of IL-6 (35), which is one of the key players of the “cytokine storm” produced by SARS-CoV-2 infection (36, 37). Lactoferrin is classified by the FDA as ‘Generally Recognised as Safe’ (GRAS) and therefore may represent a promising therapy for pre- and post-exposure prophylaxis.

Combination therapies are likely to be required for effectively treating SARS-CoV-2 infection, and this approach has already shown promise, i.e., combination therapy with interferon β-1b, lopinavir–ritonavir, and ribavirin showed efficacy against SARS-CoV-2 in a prospective, open-label, randomized, phase 2 trial (38). Here, we show that lactoferrin potentiates the antiviral activity of remdesivir and could be used in combination therapy with these drugs, which are currently being used or studied for the treatment of COVID-19. Due to its wide availability, limited cost, and strong safety profile, lactoferrin could be a rapidly deployable option for both prophylaxis and the management of COVID-19. Although our findings are promising, further studies are needed to confirm the efficacy of our lead antiviral compounds in animal models and/or clinical studies.

## METHODS

### Cells and virus

Vero E6, Caco-2, LNcaP and Huh7 cells were maintained at 37°C with 5% CO_2_ in Dulbecco’s Modified Eagle’s Medium (DMEM; Welgene), supplemented with 10% heat-inactivated fetal bovine serum (FBS), HEPES, non-essential amino-acids, L-glutamine and 1X Antibiotic-Antimycotic solution (Gibco). iPSC (SPC2 iPSC line, clone SPC2-ST-B2, Boston University) derived alveolar epithelial type 2 cells (iAEC2s) were differentiated and maintained as alveolospheres embedded in 3D Matrigel in “CK+DCI” media, as previously described (Jacob et al. 2019). iAEC2s were passaged approximately every two weeks by dissociation into single cells via the sequential application of dispase (2 mg/ml, Thermo Fisher Scientific, 17105-04) and 0.05% trypsin (Invitrogen, 25300054) and re-plated at a density of 400 cells/μl of Matrigel (Corning, 356231), as previously described (39). SARS-CoV-2 WA1 strain was obtained by BEI resources and was propagated in Vero E6 cells. Lack of genetic drift of our viral stock was confirmed by deep sequencing. Viral titers were determined by TCID50 assays in Vero E6 cells (Reed and Muench method) by microscopic scoring. All experiments using SARS-CoV-2 were performed at the University of Michigan under Biosafety Level 3 (BSL3) protocols in compliance with containment procedures in laboratories approved for use by the University of Michigan Institutional Biosafety Committee (IBC) and Environment, Health and Safety (EHS).

### Viral titer determination

Vero E6, Caco-2, LNCaP and Huh7 cells were seeded in a 48-well plate at 2×10^4^ cells/well and incubated overnight at 37°C with 5% CO_2_. Cells were then infected with SARS-CoV-2 WA1 at a multiplicity of infection (MOI) of 0.2. One hour after infection, cells were harvested (day 0 of infection) or kept at 37°C for 1, 2 and 3 days p.i.. Viral titer determination was performed by TCID50 assay on Vero E6 cells of the total virus (supernatant and intracellular fraction). Alternatively, cells were harvested with Trizol and total cellular and viral RNA was extracted with the ZymoGen Direct-zol RNA extraction kit. Viral RNA was quantified by RT-qPCR using the 2019-nCoV CDC qPCR Probe Assay and the probe set N1 (IDT technologies). IFNβ, viperin, MX1, ISG15, IFITM3 and the housekeeping gene GAPDH mRNA levels were quantified by qPCR with SsoAdvanced™ Universal SYBR® Green Supermix (Bio-Rad) with specific primers (IFNβ: F-TTGACATCCCTGAGGAGATTAAGC, R- TCCCACGTACTCCAACTTCCA; MX1: F-CCAGCTGCTGCATCCCACCC, R-AGGGGCGCACCTT CTCCTCA; ISG15: F-TGGCGGGCAACGAATT, R- GGGTGATCTGCGCCTTCA; IFITM3: F-TCCCAC GTACTCCAACTTCCA, R-AGCACCAGAAACACGTGCACT; GAPDH: F-CTCTGCTCCTCCTGTTCGAC, R-GCGCCCCACCAAGCTCAAGA). Fold increase was calculated by using the ΔΔCt method over non-infected and untreated Huh7.

### Viral infectivity assay

384-well plates (Perkin Elmer, 6057300) were seeded with Huh7 cells at 3000 cells/well and allowed to adhere overnight. Compounds were then added to the cells and incubated for 4 hours. The plates were then transferred to BSL3 containment and infected with SARS-CoV-2 WA1 at a multiplicity of infection (MOI) of 0.2 in a 10 μL addition with shaking to distribute virus. For the final dose-responses curves, porcine trypsin (Sigma-Aldrich, T0303) at a final concentration of 2 μg/ml was included during infection. After one hour of absorption, the virus inoculum was removed and fresh media with compound was added. Uninfected cells and vehicle-treated cells were included as positive and negative control, respectively. Two days post-infection, cells were fixed with 4% PFA for 30 minutes at room temperature, permeabilized with 0.3% Triton X-100, and blocked with antibody buffer (1.5% BSA, 1% goat serum and 0.0025% Tween 20). The plates were then sealed, surface decontaminated, and transferred to BSL2 for staining with the optimized fluorescent dye-set: anti-nucleocapsid protein (anti-NP) SARS-CoV-2 antibody (Antibodies Online, Cat# ABIN6952432) overnight treatment at 4 °C followed by staining with secondary antibody Alexa-647 (goat anti-mouse, Thermo Fisher, A21235), Hoechst-33342 pentahydrate (bis-benzimide) for nuclei staining (Thermo FIsher, H1398), HCS LipidTOX™ Green Neutral Lipid Stain (Thermo Fisher, H34475), and HCS CellMask™ Orange for cell delineation (Thermo Fisher H32713). iAEC2 maintained in 3D culture were dissociated to single cells and seeded in collagen coated 384-well plates at a seeding density of 8000 cells/well in the presence of 10 μM Y-27632 for the first 72 hours after plating (APExBIO, A3008) to grow to roughly 80% confluence. Infection was performed at MOI of 10 in the presence of 2 μg/ml of trypsin porcine (Sigma-Aldrich, T0303). Staining protocol for the iAEC2s differed slightly with the addition of an anti-acetylated tubulin primary antibody (Cell Signaling, 5335), instead of HCS CellMask Orange, and the use of an additional secondary Alexa 488 antibody (donkey anti-rabbit, Jackson ImmunoResearch, 711-545-152).

### Compound library

The compound library deployed for drug screening was created using the FDA-Approved Drugs Screening Library (Item No. 23538) from Cayman Chemical Company. This library of 875 compounds was supplemented with additional FDA approved drugs and selected clinical candidates from other vendors including MedChemExpress, Sigma Aldrich, and Tocris. Clinical candidates and chemical probes were included if they had any reported antiviral efficacy or had an association with SARS1, MERS or SARS-CoV-2. The library was formatted in five 384-well compound plates and was dissolved in DMSO at 10 mM. Hololactoferrin (Sigma Aldrich, L4765), apolactoferrin (Jarrow Formulas, 121011), native human lactoferrin (Creative BioMart, LFT-8196H), and transferrin (Sigma Aldrich, T2036) were handled separately and added manually in cell culture media. Dilution plates were generated for qHTS at concentrations of 2 mM, 1 mM, 500 μM, 250 μM and 50 μM and compounds were dispensed at 1:1000 dilution.

### qHTS primary screen and dose response confirmation

For the qHTS screen, compounds were added to cells using a 50 nL pin tool array on a Caliper Life Sciences Sciclone ALH 3000 Advanced Liquid Handling system. Concentrations of 2 μM, 1 μM, 500 nM, 250 nM and 50 nM were included for the primary screen. In all dose-response confirmation experiments, compounds were dispensed using an HP D300e Digital Compound Dispenser and normalized to a final DMSO concentration of 0.1% DMSO. Dose response confirmation was performed in triplicate and in 10-point:2-fold dilution. Z-primes for dose response plates ranged between 0.4-0.8.

### Imaging

Stained cell plates were imaged on both Yokogawa CQ1 and Thermo Fisher CX5 high content microscopes with a 20X/0.45NA LUCPlan FLN objective. Yokogawa CQ1 imaging was performed with four excitation laser lines (405nm/488nm/561nm/640nm) with spinning disc confocal and 100ms exposure times. Laser power was adjusted to yield optimal signal to noise ratio for each channel. Maximum intensity projection images were collected from 5 confocal planes with a 3-micron step size. Laser autofocus was performed and nine fields per well were imaged covering approximately 80% of the well area. The Thermofisher CX5 with LED excitation (386/23nm, 485/20nm, 560/25nm, 650/13nm) was also used and exposure times were optimized to maximize signal/background. Nine fields were collected at a single Z-plane as determined by image-based autofocus on the Hoechst channel. The primary qHTS screen was performed using CX5 images and all dose-response plates were imaged using the CQ1.

### Image segmentation and feature extraction and infection score

The open source CellProfiler software (12) was used in an Ubuntu Linux-based distributed Amazon AWS cloud implementation for segmentation, feature extraction, infection scoring and results were written to an Amazon RDS relational database using MySQL. A pipeline was developed to automatically identify infected cells in a field and to enable cell-level morphologic profiling. Infected areas were identified by Otsu thresholding and segmentation using the N-protein image, then all nuclei were identified in a similar manner in the Hoechst-33342 image, and the “relate objects” module was used to relate nuclei to an infected cell area. If a nucleus was found to reside within an infected area, then it and its corresponding cell area was labeled “infected”. The percentage of infected cells was tabulated by dividing the infected cell number by the total cell number summed across all 9 fields per well. To enable morphologic cell profiling, the following regions of interest were defined for feature extraction: nuclei, cell, cytoplasm, nucleoli, neutral lipid droplets and syncytia. Multiple intensity features and radial distributions were measured for each object in each channel and cell size and shape features were measured. Nuclei were segmented using the Hoechst-33342 image and the whole cell mask was generated by expanding the nuclear mask to the edge of the Cell Mask Orange image. Plate-based normalization was performed to account for variability in infection percentage. The assay window was normalized between the positive control wells (32 uninfected wells representing 0% inhibition) and the negative control wells (32 infected wells, 0.1% DMSO vehicle treated representing 100% effect). The compound treated wells were scored with the RF model and the efficacy score was normalized to the individual plate.

### Data pre-processing

Cell level data were pre-processed and analyzed in the open source Knime analytics platform (39). Cell-level data was imported into Knime from MySQL, drug treatment metadata was joined, and features were centered and scaled. Features were pruned for low variance (<5%) and high correlation (>95%) and resulted in 660 features per cell.

### Statistical methods and hypothesis testing

Dose-response curves were fit and pairwise differences between experimental conditions were tested using Prism (Graphpad Software, San Diego, CA, USA). Other statistical tests including non-parametric Mann-Whitney were performed in the statistical programming language and environment R.

### UMAP embedding

The embed_umap application of MPLearn (v0.1.0, https://github.com/momeara/MPLearn) was used to generate UMAP embeddings. Briefly, for a set of cells, each feature was per-plate standardized and jointly orthogonalized using sklearn.IncrementalPCA(n_components=379, batch_size=1000). Then, features were embedded into 2 dimensions using umap-learn (v0.4.1)(41). UMAP(n_components=2, n_neighbors=15, min_dist=0, init=‘spectral’, low_memory=True). Embeddings were visualized using Holovies Datashader (v1.12.7) (42), using histogram equalization and the viridis color map.

### Data analytics

HC Stratominer (Core Life Analytics, Utrecht NL) was used as an independent method for hit-calling and performs fully automated/streamlined cell-level data pre-processing and score generation. IC Stratominer was also used to fit dose response curves for qHTS. Compound registration and assay data registration were performed using the open source ACAS platform (Refactor BioSciences github https://github.com/RefactorBio/acas).

### Dose-response analysis and compound selection

In qHTS screening, a compound was selected to be carried forward into dose response confirmation when meeting one of the following criteria: 1) Percent infected less than 25% for the median field in at least two concentrations, 2) a dose-response relationship was observed (by inspection) across the five concentrations tested, or 3) compounds of interest not meeting this criteria were carried forward if reported positive in the literature or were being evaluated in clinical trials for COVID-19.

### Dose response analysis in the confirmation and combinatorial screening

Due to the spatial inhomogeneity of infected cells across a single well, approximately half of the fields were undersaturated, resulting in a reproducible distribution per-well. Total cell and infected cell counts were summed over the 9 fields and percent infected cells was averaged over triplicate wells. Cells treated with known fluorescent compounds, including Clofazimine, were confirmed to not have spectral interference. Dose response curves were fit with Graphpad Prism using a semilog 4-parameter variable slope model.

### Viral Binding assay

Huh-7 cells were plated in 48-well plates at 100,000 cells per well and allowed to adhere overnight. The following day, compounds were added at the indicated concentration in serum-free DMEM and incubated for 1 hour at 4 ℃. Following compound incubation, cells were infected with SARS-CoV-2 at an MOI of 10 for 1 hour at 4 ℃ to allow for viral binding. Cells were then washed 3 times with ice cold PBS to remove unbound virus and RNA was extracted by using the Direct-Zol RNA miniprep kit (Zymogen, R2052). Bound virus was then quantified by RT-qPCR (see section Viral titer determination and host gene quantification) and percentages were calculated over the infected non-treated condition.

### Multi-cycle cytopathogenic effect (CPE) reduction assay

Vero E6 were allowed to adhere overnight in 96-well cell culture plates. A 2-fold 10-point serial dilution of compounds (5000 nM-5 nM) and SARS-CoV-2 at MOI of 0.002 were added. CPE was evaluated by microscopic scoring at 5 d.p.i. The 50% inhibitory concentration (IC50) was calculated by logarithmic interpolation and is defined as the concentration at which the virus-induced CPE is reduced by 50%.

### RNAscope of SARS-CoV-2 infected cells

PFA-fixed 96-well black plates (Corning, cat nr. 3036) were permeabilized with a step-wise EtOH treatment (25% EtOH for 3 minutes, 50% EtOH for 3 minutes and 70% EtOH overnight at 4 ℃). The day after, cells were treated with washing buffer (25% formamide in 1X SSC buffer) for 5 minutes and hybridized with custom-designed probes targeting positive-sense SARS-CoV-2 RNA directly conjugated with ATTO647 (Ann Arbor Bioscience) at 37 °C overnight in hybridization buffer (dextran sulfate, 25% formamide and 0.1 % SDS). Cells were counterstained with Hoechst 33342 and anti-S protein antibody (Spike antibody 1A9, GeneTex, Cat Nr. GTX632604) and imaged using a Thermo Fisher CX5 high content microscope with a 10X/0.45NA LUCPlan FLN objective.

## Abbreviations

MOI: multiplicity of infection
COVID-19: Coronavirus Disease-2019
MOA: mechanism of action
iAEC2: induced pluripotent stem cell (iPSC)-derived alveolar epithelial type 2 cells

## ACKNOWLEDGEMENTS

Funding: University of Michigan Institute for Clinical and Health Research (MICHR) (NCATS - UL1TR002240) and its Center for Drug Repurposing. JZS is supported by the National Institute of Diabetes and Digestive and Kidney Diseases (R01DK120623). JWW is supported by the pharmacological sciences training program (PSTP) T32 training grant. CM is supported by Marie-Slodowska Curie individual fellowship (GA - 841247) and MICHR Postdoctoral Translational Scholars Program. KDA is supported by the I.M. Rosenzweig Junior Investigator Award from the Pulmonary Fibrosis Foundation. JRS is supported by the National Heart, Lung, and Blood Institute (NHLBI – R01HL119215), by the NIAID Novel Alternative Model Systems for Enteric Diseases (NAMSED) consortium (U19AI116482) and by grant number CZF2019-002440 from the Chan Zuckerberg Initiative DAF, an advised fund of Silicon Valley Community Foundation.

The authors would like to thank Matthew Chess for Amazon AWS support, Kevin Jan and Peyton Uhl at Yokogawa for imaging support, Nick Santoro at the University of Michigan Center for Chemical Genomics. We thank David Egan and Wienand Omta from Core Life Analytics for assisting high content data analytics as well as Philip Cheung and Brian Bolt at ReFactor Biosciences for assistance with HTS data registration. Finally, we thank Tracey Schultz and Dianne Jazdzyk for project management.

## SUPPLEMENTARY INFORMATION

**Supplementary Figure 1:**
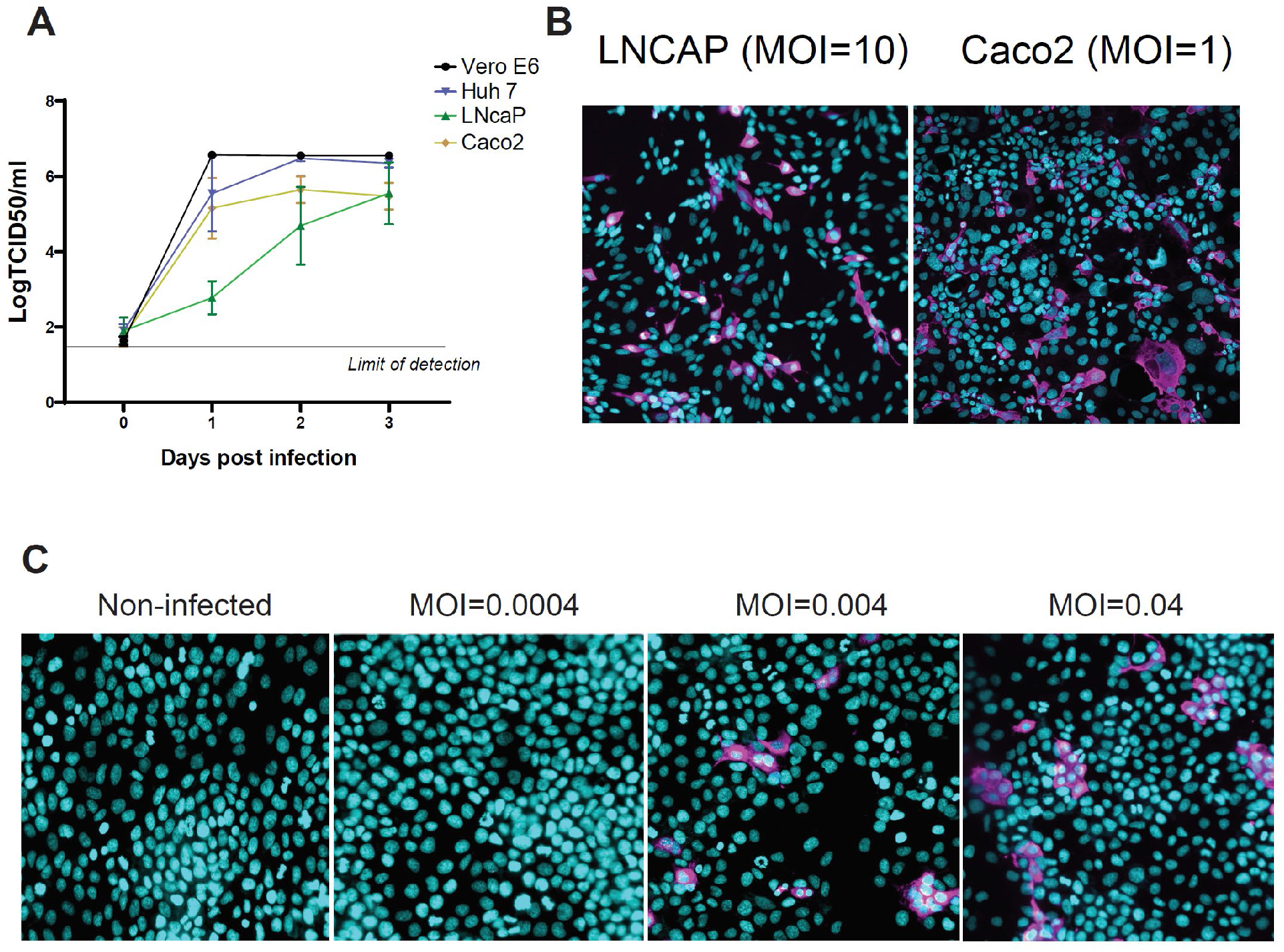
a) Growth kinetics of Vero E6, Huh-7 and Caco-2 cells. Cells were infected in 48-well plates with SARS-CoV-2 at an MOI of 0.2 and harvested at day 0 (1h post adsorption), day 1, 2 and 3. TCID_50_ determination was performed on the supernatant and cellular fraction. Graph represents median, SD of N=2 biological replicates with n=3 technical replicate each. B) Syncytia formation (magenta, anti-SARS-CoV-2 N antibody) in SARS-CoV-2 infected Vero E6 (left) and Caco2 (right) cells counter stained with Hoechst 33342 (cyan) and anti-SARS-CoV-2 N antibody (magenta).[<u>cw1</u>] C) Limit of detection of the assay. Huh-7 cells were infected at indicated MOI and imaged at 48h p.i. Progressive and more feature-rich syncytia formation was observed in correlation with an increased MOI and detection was possible with infection as low as MOI 0.004.

## REFERENCES

1. Xiao F, Tang M, Zheng X, Liu Y, Li X, Shan H. 2020. Evidence for Gastrointestinal Infection of SARS-CoV-2. Gastroenterology.

2. Lin L, Jiang X, Zhang Z, Huang S, Zhang Z, Fang Z, Gu Z, Gao L, Shi H, Mai L, Liu Y, Lin X, Lai R, Yan Z, Li X, Shan H. 2020. Gastrointestinal symptoms of 95 cases with SARS-CoV-2 infection. Gut 69:997–1001.

3. Avula A, Nalleballe K, Narula N, Sapozhnikov S, Dandu V, Toom S, Glaser A, Elsayegh D. 2020. COVID-19 presenting as stroke. Brain Behav Immun https://doi.org/10.1016/j.bbi.2020.04.077.

4. Kochi AN, Tagliari AP, Forleo GB, Fassini GM, Tondo C. 2020. Cardiac and arrhythmic complications in patients with COVID-19. J Cardiovasc Electrophysiol 31:1003–1008.

5. Mulangu S, Dodd LE, Davey RT Jr, Tshiani Mbaya O, Proschan M, Mukadi D, Lusakibanza Manzo M, Nzolo D, Tshomba Oloma A, Ibanda A, Ali R, Coulibaly S, Levine AC, Grais R, Diaz J, Lane HC, Muyembe-Tamfum J-J, PALM Writing Group, Sivahera B, Camara M, Kojan R, Walker R, Dighero-Kemp B, Cao H, Mukumbayi P, Mbala-Kingebeni P, Ahuka S, Albert S, Bonnett T, Crozier I, Duvenhage M, Proffitt C, Teitelbaum M, Moench T, Aboulhab J, Barrett K, Cahill K, Cone K, Eckes R, Hensley L, Herpin B, Higgs E, Ledgerwood J, Pierson J, Smolskis M, Sow Y, Tierney J, Sivapalasingam S, Holman W, Gettinger N, Vallée D, Nordwall J, PALM Consortium Study Team. 2019. A Randomized, Controlled Trial of Ebola Virus Disease Therapeutics. N Engl J Med 381:2293–2303.

6. Oprea TI, Bauman JE, Bologa CG, Buranda T, Chigaev A, Edwards BS, Jarvik JW, Gresham HD, Haynes MK, Hjelle B, Hromas R, Hudson L, Mackenzie DA, Muller CY, Reed JC, Simons PC, Smagley Y, Strouse J, Surviladze Z, Thompson T, Ursu O, Waller A, Wandinger-Ness A, Winter SS, Wu Y, Young SM, Larson RS, Willman C, Sklar LA. 2011. Drug Repurposing from an Academic Perspective. Drug Discov Today Ther Strateg 8:61–69.

7. McQuin C, Goodman A, Chernyshev V, Kamentsky L, Cimini BA, Karhohs KW, Doan M, Ding L, Rafelski SM, Thirstrup D, Wiegraebe W, Singh S, Becker T, Caicedo JC, Carpenter AE. 2018. CellProfiler 3.0: Next-generation image processing for biology. PLoS Biol 16:e2005970.

8. Chen W, Li X-M, Li A-L, Yang G, Hu H-N. 2016. Hepatitis C Virus Increases Free Fatty Acids Absorption and Promotes its Replication Via Down-Regulating GADD45α Expression. Med Sci Monit 22:2347–2356.

9. Riva L, Yuan S, Yin X, Martin-Sancho L, Matsunaga N, Pache L, Burgstaller-Muehlbacher S, De Jesus PD, Teriete P, Hull MV, Chang MW, Chan JF-W, Cao J, Poon VK-M, Herbert KM, Cheng K, Nguyen T-TH, Rubanov A, Pu Y, Nguyen C, Choi A, Rathnasinghe R, Schotsaert M, Miorin L, Dejosez M, Zwaka TP, Sit K-Y, Martinez-Sobrido L, Liu W-C, White KM, Chapman ME, Lendy EK, Glynne RJ, Albrecht R, Ruppin E, Mesecar AD, Johnson JR, Benner C, Sun R, Schultz PG, Su AI, García-Sastre A, Chatterjee AK, Yuen K-Y, Chanda SK. 2020. Discovery of SARS-CoV-2 antiviral drugs through large-scale compound repurposing. Nature 586:113–119.

10. Heiser K, McLean PF, Davis CT, Fogelson B, Gordon HB, Jacobson P, Hurst B, Miller B, Alfa RW, Earnshaw BA, Victors ML, Chong YT, Haque IS, Low AS, Gibson CC. 2020. Identification of potential treatments for COVID-19 through artificial intelligence-enabled phenomic analysis of human cells infected with SARS-CoV-2. bioRxiv.

11. Jeon S, Ko M, Lee J, Choi I, Byun SY, Park S, Shum D, Kim S. Identification of antiviral drug candidates against SARS-CoV-2 from FDA-approved drugs.

12. Yang L, Pei R-J, Li H, Ma X-N, Zhou Y, Zhu F-H, He P-L, Tang W, Zhang Y-C, Xiong J, Xiao S-Q, Tong X-K, Zhang B, Zuo J-P. 2020. Identification of SARS-CoV-2 entry inhibitors among already approved drugs. Acta Pharmacol Sin https://doi.org/10.1038/s41401-020-00556-6.

13. Hurley K, Ding J, Villacorta-Martin C, Herriges MJ, Jacob A, Vedaie M, Alysandratos KD, Sun YL, Lin C, Werder RB, Huang J, Wilson AA, Mithal A, Mostoslavsky G, Oglesby I, Caballero IS, Guttentag SH, Ahangari F, Kaminski N, Rodriguez-Fraticelli A, Camargo F, Bar-Joseph Z, Kotton DN. 2020. Reconstructed Single-Cell Fate Trajectories Define Lineage Plasticity Windows during Differentiation of Human PSC-Derived Distal Lung Progenitors. Cell Stem Cell 26:593–608.e8.

14. Mason RJ. 2020. Pathogenesis of COVID-19 from a cell biology perspective. Eur Respir J 55.

15. Lang J, Yang N, Deng J, Liu K, Yang P, Zhang G, Jiang C. 2011. Inhibition of SARS pseudovirus cell entry by lactoferrin binding to heparan sulfate proteoglycans. PLoS One 6:e23710.

16. Kell DB, Heyden EL, Pretorius E. 2020. The Biology of Lactoferrin, an Iron-Binding Protein That Can Help Defend Against Viruses and Bacteria. Front Immunol 11:1221.

17. Baeuerle PA, Huttner WB. 1986. Chlorate — a potent inhibitor of protein sulfation in intact cells. Biochemical and Biophysical Research Communications.

18. Siqueiros-Cendón T, Arévalo-Gallegos S, Iglesias-Figueroa BF, García-Montoya IA, Salazar-Martínez J, Rascón-Cruz Q. 2014. Immunomodulatory effects of lactoferrin. Acta Pharmacol Sin 35:557–566.

19. Blanco-Melo D, Nilsson-Payant BE, Liu W-C, Uhl S, Hoagland D, Møller R, Jordan TX, Oishi K, Panis M, Sachs D, Wang TT, Schwartz RE, Lim JK, Albrecht RA, tenOever BR. 2020. Imbalanced Host Response to SARS-CoV-2 Drives Development of COVID-19. Cell 181:1036–1045.e9.

20. Kuleshov MV, Stein DJ, Clarke DJB, Kropiwnicki E, Jagodnik KM, Bartal A, Evangelista JE, Hom J, Cheng M, Bailey A, Zhou A, Ferguson LB, Lachmann A, Ma’ayan A. 2020. The COVID-19 Drug and Gene Set Library. Patterns.

21. Yin W, Mao C, Luan X, Shen D-D, Shen Q, Su H, Wang X, Zhou F, Zhao W, Gao M, Chang S, Xie Y-C, Tian G, Jiang H-W, Tao S-C, Shen J, Jiang Y, Jiang H, Xu Y, Zhang S, Zhang Y, Xu HE. 2020. Structural basis for inhibition of the RNA-dependent RNA polymerase from SARS-CoV-2 by remdesivir. Science 368:1499–1504.

22. Ghahremanpour MM, Tirado-Rives J, Deshmukh M, Ippolito JA, Zhang C-H, de Vaca IC, Liosi M-E, Anderson KS, Jorgensen WL. Identification of 14 Known Drugs as Inhibitors of the Main Protease of SARS-CoV-2.

23. Roscow O, Ganassin R, Garver K, Polinski M. 2018. Z-FA-FMK demonstrates differential inhibition of aquatic orthoreovirus (PRV), aquareovirus (CSRV), and rhabdovirus (IHNV) replication. Virus Res 244:194–198.

24. Ou X, Liu Y, Lei X, Li P, Mi D, Ren L, Guo L, Guo R, Chen T, Hu J, Xiang Z, Mu Z, Chen X, Chen J, Hu K, Jin Q, Wang J, Qian Z. 2020. Characterization of spike glycoprotein of SARS-CoV-2 on virus entry and its immune cross-reactivity with SARS-CoV. Nat Commun 11:1620.

25. Pardanani A, Hood J, Lasho T, Levine RL, Martin MB, Noronha G, Finke C, Mak CC, Mesa R, Zhu H, Others. 2007. TG101209, a small molecule JAK2-selective kinase inhibitor potently inhibits myeloproliferative disorder-associated JAK2 V617F and MPL W515L/K mutations. Leukemia 21:1658–1668.

26. Wu D, Yang XO. 2020. TH17 responses in cytokine storm of COVID-19: An emerging target of JAK2 inhibitor Fedratinib. J Microbiol Immunol Infect https://doi.org/10.1016/j.jmii.2020.03.005.

27. Zhang W, Zhao Y, Zhang F, Wang Q, Li T, Liu Z, Wang J, Qin Y, Zhang X, Yan X, Zeng X, Zhang S. 2020. The use of anti-inflammatory drugs in the treatment of people with severe coronavirus disease 2019 (COVID-19): The Perspectives of clinical immunologists from China. Clinical Immunology.

28. Stebbing J, Phelan A, Griffin I, Tucker C, Oechsle O, Smith D, Richardson P. 2020. COVID-19: combining antiviral and anti-inflammatory treatments. Lancet Infect Dis. thelancet.com.

29. Treatment of Moderate to Severe Coronavirus Disease (COVID-19) in Hospitalized Patients - Full Text View - ClinicalTrials.gov.

30. Cai Y, Liu Y, Zhang X. 2007. Suppression of coronavirus replication by inhibition of the MEK signaling pathway. J Virol 81:446–456.

31. Gu J, Han B, Wang J. 2020. COVID-19: gastrointestinal manifestations and potential fecal--oral transmission. Gastroenterology 158:1518–1519.

32. Reghunathan R, Jayapal M, Hsu L-Y, Chng H-H, Tai D, Leung BP, Melendez AJ. 2005. Expression profile of immune response genes in patients with Severe Acute Respiratory Syndrome. BMC Immunol 6:2.

33. Han C, Duan C, Zhang S, Spiegel B, Shi H, Wang W, Zhang L, Lin R, Liu J, Ding Z, Hou X. 2020. Digestive Symptoms in COVID-19 Patients With Mild Disease Severity: Clinical Presentation, Stool Viral RNA Testing, and Outcomes. Am J Gastroenterol https://doi.org/10.14309/ajg.0000000000000664.

34. Oda H, Kolawole AO, Mirabelli C, Wakabayashi H, Tanaka M, Yamauchi K, Abe F, Wobus CE. 2020. Antiviral Effects of Bovine Lactoferrin on Human Norovirus. Biochem Cell Biol https://doi.org/10.1139/bcb-2020-0035.

35. Cutone A, Frioni A, Berlutti F, Valenti P, Musci G, Bonaccorsi di Patti MC. 2014. Lactoferrin prevents LPS-induced decrease of the iron exporter ferroportin in human monocytes/macrophages. Biometals 27:807–813.

36. Conti P, Ronconi G, Caraffa A, Gallenga C, Ross R, Frydas I, Kritas S. 2020. Induction of pro-inflammatory cytokines (IL-1 and IL-6) and lung inflammation by Coronavirus-19 (COVI-19 or SARS-CoV-2): anti-inflammatory strategies. J Biol Regul Homeost Agents 34.

37. Lagunas-Rangel FA, Chávez-Valencia V. 2020. High IL-6/IFN-γ ratio could be associated with severe disease in COVID-19 patients. J Med Virol https://doi.org/10.1002/jmv.25900.

38. Hung IF-N, Lung K-C, Tso EY-K, Liu R, Chung TW-H, Chu M-Y, Ng Y-Y, Lo J, Chan J, Tam AR, Shum H-P, Chan V, Wu AK-L, Sin K-M, Leung W-S, Law W-L, Lung DC, Sin S, Yeung P, Yip CC-Y, Zhang RR, Fung AY-F, Yan EY-W, Leung K-H, Ip JD, Chu AW-H, Chan W-M, Ng AC-K, Lee R, Fung K, Yeung A, Wu T-C, Chan JW-M, Yan W-W, Chan W-M, Chan JF-W, Lie AK-W, Tsang OT-Y, Cheng VC-C, Que T-L, Lau C-S, Chan K-H, To KK-W, Yuen K-Y. 2020. Triple combination of interferon beta-1b, lopinavir–ritonavir, and ribavirin in the treatment of patients admitted to hospital with COVID-19: an open-label, randomised, phase 2 trial. Lancet https://doi.org/10.1016/S0140-6736(20)31042-4.

39. Jacob A, Morley M, Hawkins F, McCauley KB, Jean JC, Heins H, Na C-L, Weaver TE, Vedaie M, Hurley K, Hinds A, Russo SJ, Kook S, Zacharias W, Ochs M, Traber K, Quinton LJ, Crane A, Davis BR, White FV, Wambach J, Whitsett JA, Cole FS, Morrisey EE, Guttentag SH, Beers MF, Kotton DN. 2017. Differentiation of Human Pluripotent Stem Cells into Functional Lung Alveolar Epithelial Cells. Cell Stem Cell 21:472–488.e10.

40. Berthold MR, Cebron N, Dill F, Gabriel TR, Kötter T, Meinl T, Ohl P, Thiel K, Wiswedel B. 2009. KNIME - the Konstanz information miner. ACM SIGKDD Explorations Newsletter.

41. McInnes L, Healy J, Melville J. 2018. UMAP: Uniform Manifold Approximation and Projection for Dimension Reduction. arXiv [statML].

42. Stevens J-LR, Rudiger P, Bednar JA. 2015. HoloViews: Building complex visualizations easily for reproducible science, p. 61–69. In Proceedings of the 14th Python in Science Conference.

